# A population of CD4^+^ T cells with a naïve phenotype stably polarized to the T_H_1 lineage

**DOI:** 10.1101/2020.07.14.202168

**Authors:** Jonathan W. Lo, Maria Vila de Mucha, Luke B. Roberts, Natividad Garrido-Mesa, Arnulf Hertweck, Joana F. Neves, Emilie Stolarczyk, Stephen Henderson, Ian Jackson, Jane K. Howard, Richard G. Jenner, Graham M. Lord

## Abstract

T-bet is the lineage-specifying transcription factor for CD4^+^ T helper type 1 (T_H_1) cells. T-bet has also been found in other CD4^+^ T cell subsets, including T_H_17 cells and T_REG_, where it modulates their functional characteristics. However, we lack information on when and where T-bet is expressed during T cell differentiation and how this impacts T cell function. To address this, we traced the ontogeny of T-bet-expressing cells using a fluorescent fate-mapping mouse line. We demonstrate that T-bet is expressed in a subset of CD4^+^ T cells with naïve cell surface markers and that this novel cell population is phenotypically and functionally distinct from conventional naïve CD4^+^ T cells. These cells are also distinct from previously described populations of memory phenotype and stem cell-like T cells. Naïve-like T-bet-experienced cells are polarised to the T_H_1 lineage, predisposed to produce IFNγ upon cell activation, and resist repolarisation to other lineages *in vitro* and *in vivo*. These results demonstrate that lineage-specifying factors can function to polarise T cells in the absence of canonical markers of T cell activation and that this has an impact on the subsequent T helper response.

## Introduction

Upon encounter with specific antigen, naïve CD4^+^ T cells differentiate into one of several T helper cell subtypes, including T_H_1, T_H_2, T_H_17, follicular T cells (T_FH_) and peripheral regulatory T cells (pT_REG_). CD4^+^ T cell fate depends on the cytokine environment, signalling through the T cell receptor (TCR), and the transcription factors that these induce. Differentiation of T_H_1 cells is triggered in response to presentation of bacterial antigens by antigen presenting cells (APCs) (1) and requires IL-12-mediated activation of STAT4 (2). T_H_1 cells are characterised by their production of interferon gamma (IFNγ), which activates cell-mediated immunity against intracellular bacteria, viruses, and tumour cells. Inappropriate or excessive T_H_1 responses also contribute to inflammatory diseases, highlighting the importance of understanding the regulation of T_H_1 cell differentiation.

Differentiation of naïve T cells into T_H_1 effectors requires the T-box transcription factor T-bet, encoded by *Tbx21*, which is upregulated via IL12-dependent activation of STAT4 (2, 3). T-bet directly activates *Ifng*, and a number of other genes encoding cytokines and receptors, including *Il12rb2*, *Cxcr3* and *Ccl4* (2, 4–6). T-bet also activates its own expression, by both directly binding to its own gene and via IFNγ-mediated activation of STAT1 (7, 8). T-bet mediated *Ifng* activation is accompanied by chromatin modifications that are maintained in memory cells (9–11). In addition to promoting T_H_1 lineage-specification, T-bet also counteracts differentiation of CD4^+^ T cells into other lineages. T-bet represses expression of the gene encoding the T_H_2 lineage-specifying factor *Gata3* (12) and blocks GATA3 binding to its target genes (8, 13). T-bet also supresses T_H_17 differentiation by blocking upregulation of the genes encoding RORγt and IRF4 (14, 15).

Although serving as the T_H_1 lineage-specifying transcription factor, under certain conditions T-bet can also be expressed in T_H_2, T_H_17, pT_REG_ and in a T_FH_-T_H_1 transitional state, revealing a level of plasticity in CD4^+^ T cell differentiation (16–18). Functionally, T-bet expression in T_H_17 cells is associated with disease in murine allergic encephalomyelitis (19–21) and T-bet is required for T_REG_ homoeostasis and function during T_H_1-mediated inflammation (22). T-bet also plays key roles outside of CD4^+^ T cells, including CD8^+^ cytotoxic T cells, γδ T cells, natural killer (NK) cells, natural killer T (NKT) cells, type I and type 3 innate lymphoid cells (ILC1s and ILC3s), dendritic cells (DCs), monocytes and B cells (3).

Dysregulated T-bet function is associated with inflammatory disease. T-bet is upregulated in lamina propria T cells of patients with Crohn’s and celiac disease and *ex vivo* culture of biopsies from untreated celiac patients with gliadin peptides increases T-bet expression through STAT1 activation (23, 24). Genetic variants associated with ulcerative colitis and Crohn’s disease are enriched at T-bet binding sites and can alter T-bet binding to its target genes (25). Transfer of *Tbx21*^*−/−*^ naïve CD4^+^ T cells into *Rag2*^*−/−*^ mice gave rise to a higher proportion of IL-17A-producing CD4^+^ T cells and worse colitis compared to mice which received WT naïve CD4^+^ T cells (15, 26), suggesting that T-bet restrains pathology during IL-17A-driven colitis (26–29).

Naïve T cells can be identified by their surface markers and can be defined as CD62L^high^ CD44^low^ CCR7^high^ T cells. Consistent with their quiescent, non-antigen experienced state, naïve CD4^+^ T cells have not been found to express lineage-specifying transcription factors, apart from GATA3, which is essential for CD4^+^ T cell homeostasis in addition to its role in T_H_2 cell differentiation. In contrast, antigen-experienced central memory T cells (T_CM_) and effector memory T cells (T_EM_) are defined as CD62L^high^ CD44^high^ CCR7^high^ and CD62L^low^ CD44^high^ CCR7^low^, respectively (30–33). However, the distinction between the naïve and memory cell states is not always clear cut. Small numbers of memory-phenotype (MP) cells are found in normal, non-immunised mice (34–36). MP cells, which are predominantly CD8^+^, also arise in mice maintained under germ-free and antigen-free conditions and in humans before birth, indicating they develop in response to self-antigens (37, 38). Within the MP cell population, virtual memory (VM) T cells (CD8^+^ CD62L^low^ CD44^high^ CD122^high^ CXCR3^high^ Ly6C^high^ CD49d^low^) are highly proliferative and provide both antigen-specific immunity and exhibit bystander killing activity (37, 39, 40). Memory T cells with a naïve phenotype (T_MNP_) are a human CD8^+^ T cell population defined as CXCR3^+^ CD49d^+^ CCR7^+^ CD95^lo^ CD28^int^ and are also highly responsive to virus peptides (41). T_H_1-like memory phenotype (T_H_1-like MP) T cells (CD4^+^ CD62L^−^CD44+ CXCR3^+^ ICOS^+^ CD49d^high^) express high levels of T-bet and are rapidly able to produce T_H_1 cytokines, like IFNγ, without the need for TCR activation (42). Like naïve T cells, CD8^+^ T memory stem cells (T_SCM_) are CD62L^high^ CD44^low^ and exhibit high proliferative capacity, but co-express markers CD122 (IL-2Rβ), CXCR3, Sca-1, Bcl2 and low levels of T-bet (43–45). A population of MP CD4^+^ cells has also been identified (46, 47). These cells are present in germ-free and antigen-experienced mice and develop from CD5^hi^ naïve cells in the periphery in a TCR and CD28-dependent manner (46). MP cells contain T-bet^lo^, T-bet^int^ and T-bet^hi^ subpopulations, with T-bet^hi^ MP cells providing a rapid, non-antigen specific, upregulation of IFNγ in response to infection (46).

Knowledge of the CD4^+^ T cell subsets that express T-bet, the points during T cell ontogeny at which T-bet is expressed, and the impact of T-bet on cell function is important for understanding T cell differentiation and its dysregulation in disease. However, progress has been limited by our inability to identify cells that have experienced T-bet expression. To address this, we have developed a T-bet^cre^ × Rosa26-YFP^fl-STOP-fl^ mouse line in which YFP marks cells that have expressed T-bet during their ontogeny. Using this line, we report the discovery of a previously unidentified CD4^+^ T cell population with a naïve cell surface phenotype, which, nevertheless, has experienced T-bet expression. Arising shortly after birth, these YFP^+^ naïve CD4^+^ T cells accounted for around 1% of splenic CD4^+^ T cells and are distinct from previously identified populations of memory-like cells. Although displaying naïve cell surface markers, these T-bet-experienced cells were polarised towards the T_H_1 lineage, predisposed to rapidly upregulate IFNγ upon activation, and stably maintained their phenotype in opposing polarisation conditions in vitro and in an *in vivo* colitis model. This work reveals that naïve-like CD4^+^ T cells can express lineage-specific transcription factors and that this shapes subsequent immune responses.

## Materials and Methods

### Animals

C56BL/6 *Rag1*^*−/−*^ (Jackson labs) and *Rosa26*^*YFP/+*^ (Jackson labs) mice were sourced commercially. *T-bet*^*cre/+*^ mice were previously generated by our group. *T-bet*^*cre*/+^ mice were bred with *Rosa26*^*YFP/+*^ to generate the *T-bet*^*cre/+*^ *Rosa26*^*YFP/+*^ line (hereby referred to as T-bet^FM^ mice). The *T-bet*^*Amcyan*^ line was a kind gift obtained from Jinfang Zhu at NIH/NIAID (National Institute of Health-National Institute of Allergy and Infectious Diseases) in Maryland, USA. All mice used were aged between 6-12 weeks, unless stated otherwise. Foetal mice were culled at E.15.5 in accordance with humane Schedule 1 practice set out by the Home Office. All animal experiments were performed in accredited facilities in accordance with the UK Animals (Scientific Procedures) Act 1986 (Home Office Licence Numbers PPL: 70/6792 and 70/8127).

### Cell isolation and preparation

Adult mice were euthanized using an approved Schedule 1 method of inhalation of a rising concentration of carbon dioxide gas followed by cervical dislocation. Embryo and neonatal mice were euthanised by using an approved Schedule 1 method of decapitation. Spleen, thymus, liver, mesenteric lymph nodes (mLN), peripheral (axillary, inguinal and cervical) lymph nodes (pLN) and colons were excised and placed in cold phosphate buffered saline (PBS) solution.

Colonic lamina propria cells were isolated as described (48). Colons were cleaned and fat and faeces were removed. Afterwards, they were cut into 1-2 cm pieces using surgical scissors and put into 10 ml of Hank’s balanced salt solution (HBSS) without Mg^2+^/Ca^2+^ (Invitrogen) mixed with 5 mM EDTA and 10 mM HEPES (Fisher Scientific) and incubated at 37.5°C with agitation for 20 mins. Next, intestinal pieces were filtered, and the subsequent intestinal pieces were sliced into fine pieces using scalpels and were collected in batch tested complete animal medium (RPMI (Gibco) with 10% heat-inactivated foetal calf serum (FCS) (Gibco,), 2 mM glutamine, 100 U/ml penicillin and 100 µg/ml streptomycin, HEPES (Fisher Scientific), non-essential amino acids, sodium pyruvate and 2-mercaptoethanol (Sigma)). Collagenase (0.5 mg/ml, Roche), DNase (10 μg/ml, Roche) and dispase II (1.5 mg/ml, Roche) were added and the intestinal pieces incubated for a further 20 mins at 37°C with agitation. After incubation, the digestion mix was filtered once more and centrifuged at 860g for 10 mins at 4°C. Intestinal lamina propria cells were collected at the interface after centrifugation through Percol and washed.

Splenic, thymus, liver, mLN and pLN cells were isolated into a single cell suspension in complete animal medium with the use of 70 µM mesh filter and general mechanical destruction. The suspension was centrifuged at 860g for 5 mins at 4°C and mLN and pLN. spleen, thymus and liver cell pellets resuspended, and red blood cells were lysed using a standard red blood lysis buffer (ACK). The liver pellet was then collected at the interface after centrifugation through Percol and washed.

### Cell sorting

CD4^+^ cells from the spleen of mice were purified using LS positive selection magnetic-activated columns (MACs) and anti-CD4 (L3T4) beads (Miltenyi Biotec). CD4^+^ MACs sorted cells were further purified by fluorescence activated cell sorting using a FACS Aria (BD Biosciences) with a 70µm nozzle insert and FACS Diva software CD4^+^ MACs sorted cells were stained for 20 mins at 4°C in the dark with the following antibodies: anti-CD4-PerCPCy5.5 (RM4-5), anti-CD25-PE (PC61), anti-CD62L-PECy7 (MEL-14; Thermo Fisher) and anti-CD44-Pacific Blue (IM1.8.1; Thermo Fisher). Single positive compensation controls and unstained controls were used to set up instrument settings and for gating strategies. Cell purity was verified post-sort (requiring 95% purify) and cell viability assessed using trypan-blue staining. Sorted cells were collected in complete media (as described above) and after cells were counted using a haemocytometer they were immediately cultured or transferred *in vivo*.

### Naïve T cell transfer model of colitis

Naïve T cells were transferred as described (28). Spleens and mLNs were harvested from either WT C56/BL6 mice or T-bet^FM^ donor mice and mechanically disrupted, as described before. Naïve CD4^+^ T cells (CD4^+^ CD25^−^CD44^low^ CD62L^high^) were sorted using a FACS Aria to a purity of >95%, washed and resuspended in sterile PBS. *Rag2*^*−/−*^ mice were injected intraperitoneally with 5 × 105 naïve CD4^+^ cells per mouse, unless stated otherwise, and humanely culled after 6-8 weeks. Mice were monitored for their health every week for signs of illness.

### Cell culture

Unfractionated single cell suspensions of splenocytes (2×10^6^/ml), mLN (1×10^6^/ml) and cLP cells (1×10^6^/ml) were cultured in complete animal medium (same as described above) for 48 hours in a CO_2_ controlled incubator at 37°C and 5% CO_2_.

In accordance with MIATA, sorted CD4^+^ T cells from the spleens of the T-bet^FM^ mouse (as described above) were cultured at a concentration of 1 million cells per ml in complete media on pre-incubated anti-CD3/anti-CD28 bound plates in IL-2 (20ng/ml) (Sigma-Aldrich) for 2 days and then just IL-2 (20ng/ml) (Sigma-Aldrich) for a further 5 days. Supernatants were harvested and cytokine concentrations measured by ELISA (Thermo Fisher) and cells were stained and analysed using flow cytometry as described below.

### CD4^+^ T cell polarisation

Sorted CD4^+^ T cells from the spleens of the T-bet^FM^ mouse (as described above) were cultured at a concentration of 1 million cells per ml in complete media on anti-CD3/anti-CD28-bound plates for 2 days and specific cytokines added for either T_H_0: (hIL-2 (20ng/ml)), T_H_1: (anti-IL-4 (5µg/ml), mIL-12 (20ng/ml), hIL-2 (20ng/ml)), T_H_2: (anti-IFNγ (20ng/ml), mIL-4 (20ng/ml), hIL-2 (20ng/ml)), T_H_17: (anti-CD28 (5µg/ml), anti-IL-4 (5µg/ml), anti-IFNγ (20ng/ml), mIL-1β (10ng/ml), mIL-6 (20ng/ml), hTGFβ (2ng/ml)) and T_REG_: (hTGFβ (2ng/ml), hIL-2 (20ng/ml)) polarising conditions for a further 5 days. Supernatants were harvested and cytokine concentrations measured by ELISA (Thermo Fisher) and cells were stained and analysed using flow cytometry as described below.

### *Ex vivo* colon organ culture

3 mm punch biopsies (Miltex) were acquired from murine colons at full thickness. 3 biopsies were cultured in 500 μl of complete animal RPMI for 48 hours in a CO_2_ controlled incubator at 37°C and 5% CO_2_. Culture supernatants were harvested, and cytokine concentrations measured by ELISA (Thermo Fisher).

### ELISA

Cytokine concentrations for IL-17A and IFNγ from the supernatants of cultured cells and ex vivo colon organ cultures were measured by ELISA (Thermo Fisher Ready-Set-Go). Samples and standards were measured in duplicate. Standard curves were created with the standards provided in the kits, in accordance with the manufacturer’s protocols.

### Flow cytometry

Single cell suspensions extracted from the various tissues were plated out into flow cytometry tubes (Sarstedt) at a concentration of 10^6^ cells per ml. Cells were stimulated with 50 ng/ml phorbol 12-myristate 13-acetate (PMA) (Sigma Aldrich) and 1 μg/ml ionomycin (Sigma Aldrich) for 4 hours. For intracellular staining, 2 μM monensin (Sigma Aldrich) was added for the final 2 hours. For intracellular staining, cells were fixed and permeabilised using the Foxp3 fixation/permeabilization buffer kit (BD Biosciences). FcR receptor blocking antibodies were added for 15 min at 4°C and surface staining antibodies added together with live/dead stain (Fixable Live/Dead stains Thermo Fisher) and incubated for 20 mins at room temperature in the dark. Cells were acquired within 24 hours of staining. Fluorochromes were purchased from either BD Biosciences, Biolegend or Thermo Fisher. Fluorochromes used were CD4 (RM4-5), CD8α (53-6.7), CD45.2 (104) CD45.1 (A20), CD3 (17A2), CD5 (53-7.3), CD44 (IM7), CD62L (MEL-14), CD25 (PC61), CD127 (A7R34), CD27 (LG.7F9), CD28 (37.51), CD49d (R1-2), CD95 (15A7), CD11a (M17/4), CD122 (TM-b1), CCR7 (4B12), CXCR3 (CXCR3-173), T-bet (4B10), RORγt (B2D), Foxp3 (FJK-16S), GATA3 (LS0-823), IL-10 (JEs5-16E3), IL-5 (TRFK5), IL-13 (eBio13A), IFNγ (XMG1.2), γδTCR (GL3), CD1d Tetramer (PBS57-loaded or - unloaded, provided by NIH Tetramer Core Facility), B220 (RA3-6B2), DX5 (DX5), CD11c (N418), CD11b (M1/70), Nkp46 (29A1.4), NK1.1 (PK136), c-kit (ACK2), Sca-1 (D7), Haemotopoietic Lineage Cocktail (consisting of: CD3 (17A2), B220 (RA3-6B2), CD11b (M1/70), TER-119 (TER-119), Gr-1 (RB6-8C5)), ICOS (C398.4A). After incubation, cells were washed in sterile PBS and analysed with a LSRFortessa machine (BD Biosciences).

### RNA sequencing (RNA-seq)

Naïve and effector memory CD4^+^ T cells were sorted using the protocol, as described above using anti-CD4-PerCPCy5.5 (RM4-5), anti-CD25-PE (PC61), anti-CD62L-PECy7 (MEL-14; Thermo Fisher) and anti-CD44-Pacific Blue (IM1.8.1; Thermo Fisher). YFP^+^ and YFP^−^ naïve and memory cells were sorted on their basis of YFP expression. Cells were sorted into 1 ml of lysis buffer and either immediately processed for RNA extraction or frozen at −80 for RNA extraction later using the RNAeasy Micro kit (Qiagen). RNA samples were then checked for the quality, contamination and concentration using a NanoDrop, Qubit spectrophotometer and Agilent Bioanalyzer. Libraries were prepared using the Ovation SoLo RNA-seq library preparation kit (NuGEN genomics) according to the manufacturer’s protocol and were sequenced on an Illumina HiSeq 3000 to generate 2×75 bp paired-end reads. Reads were filtered to remove adaptors and low-quality bases and aligned to mm9 using TopHat2 (49). PCR duplicates were removed using the NuGEN duplicate marking tool and read counts generated using featureCounts (50). Differential gene expression analysis between YFP^+^ and YFP^−^ naïve or YFP^+^ and YFP-T_EM_ CD4^+^ T cells was conducted using DEseq2 (51), and gene expression levels across all samples were calculated and normalised using the regularized logarithm transformation (51). GSEA was performed using the fgsea package using 1000 permutations (52) with a T_H_1 gene signature identified from comparison between 3 different T_H_1 vs T_H_2 gene expression datasets (53–55). Fastq files were downloaded from the SRA and gene centred expression estimates made using kallisto together with the Gencode M20 transcript models. T_H_1-specific genes were then identified using DESeq2, with study and cell type (T_H_1/T_H_2) treated as covariates for batch correction.

### Statistical analysis

Statistical analyses were carried using GraphPad Prism 8 (GraphPad Software Inc.). Significance was calculated using the Mann-Whitney test or, for grouped data, the 1-way ANOVA test with Dunn’s corrections. Alpha was set at 0.05.

## Results

### Identification of a population of CD4^+^ T cells with a naïve surface phenotype that have previously expressed T-bet

We generated a fluorescent T-bet fate mapping (T-bet^FM^) mouse line to identify cells that have expressed T-bet during their ontogeny. We first inserted an IRES-Cre cassette in exon 6 of the endogenous locus *Tbx21* downstream of the stop codon (Supplemental Fig. 1A). These *T-bet*^*cre/+*^ mice were then crossed with *Rosa26*^*YFPfl/+*^ to generate a T-bet^FM^ line in which induction of *Tbx21* triggers *Cre* expression, removal of the *Eyfp* stop codon and thus YFP production (Supplemental Fig. 1B). Thus, this T-bet^FM^ line allows identification of cells that are expressing, or that have ever previously expressed, T-bet (Supplemental Figs. 1C and D).

We first sought to validate our model by characterising YFP expression in a basic immunophenotyping strategy for cell types that are known to express T-bet (Supplemental Figs. 1E-G). We found that 11% of CD4^+^ T cells and 20% of CD8^+^ T cells from the spleens of these mice were YFP^+^. The proportions YFP^+^ of F4/80^+^, CD11c^+^ and CD19^+^ cells in the spleen were relatively low (6%, 11% and 3%, respectively; Figs. 1A and B), consistent with the previously characterised expression of T-bet in subsets of these cell types (3). In contrast, 96% of splenic CD45^+^ NKp46^+^ lineage negative and CD45^+^ NKp46^+^ lineage positive cells were YFP^+^ (Figs. 1A and B), consistent with the high levels of T-bet expression known to be exhibited by these cell types (3).

**Figure 1.**
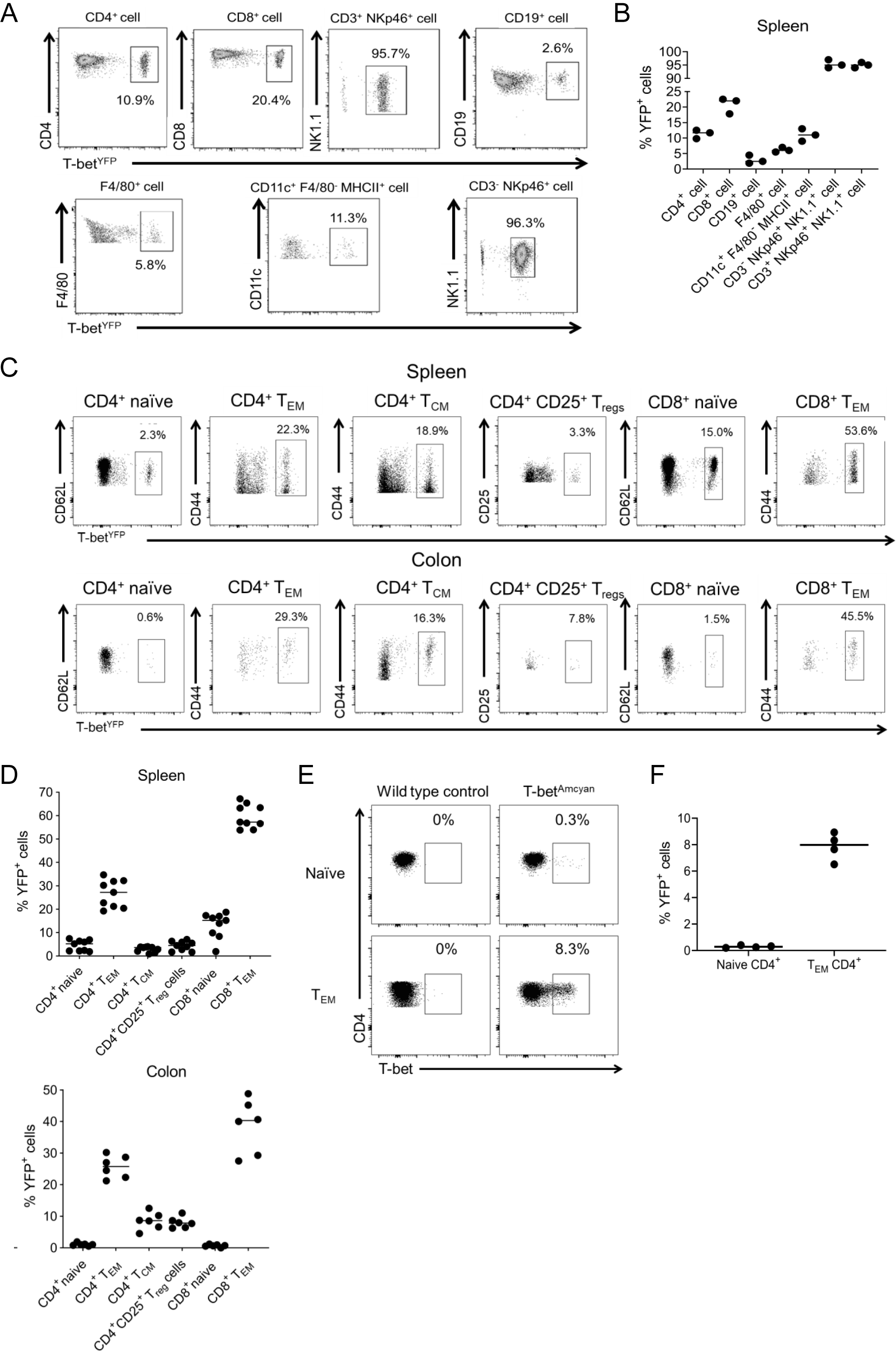
T-bet fate mapping identifies a population of peripheral naïve-like CD4^+^ T cells that have experienced T-bet expression. A. Representative flow plots showing YFP expression in different splenic cells populations. B. Quantification of the proportions of YFP+ cells for each cell type shown in A (n=3). C. Representative flow plots showing YFP expression in naïve CD4^+^ T cells (CD3^+^ CD4^+^ CD8^−^ CD25^−^ CD62L^+^ CD44^−^), effector CD4^+^ T cells (CD3^+^ CD4^+^ CD8^−^ CD25^−^ CD62L^−^ CD44^+^), regulatory T cells (CD3^+^ CD4^+^ CD8^−^ CD25^+^), naïve CD8^+^ T cells (CD3^+^ CD4^−^ CD8^+^ CD62L^+^ CD44^−^), effector CD8^+^ T cells (CD3^+^ CD4-CD8^+^ CD62L^−^ CD44^+^) and central memory CD8^+^ T cells (CD3^+^ CD4^−^ CD8^+^ CD62L^+^ CD44^+^) from the spleen (above) and colon (below). D. Mean proportions of YFP^+^ cells from the different subtypes of T cells in the spleen (n=9) and colon shown in in E (n=6). E. Representative flow plot showing the proportion of CD3^+^ CD4^+^ naïve or T_EM_ cells expressing AmCyan in the T-betAmCyan mouse reporter line. F. Overall dot plot showing the proportion of CD3^+^ CD4^+^ naïve or T_EM_ cells expressing AmCyan in the T-bet^AmCyan^ mouse reporter line (n=4).

We sought to further understand the nature of T-bet expression in CD4^+^ and CD8^+^ T cells in spleen and colon. Around a quarter of CD4^+^ T_EM_ were YFP^+^ in both the spleen and the colon (Figs. 1C and 1D). Some T_REG_ populations have previously been found to express T-bet (22, 56), and, consistent with this, around 6% of splenic and colonic CD4^+^ CD25^+^ cells expressed YFP (Figs. 1C and 1D). For CD8^+^ T_EM_ cells, this proportion of YFP^+^ cells increased to ~60% in the spleen and 40% in the colon (Figs. 1C and D).

T-bet expression is thought to be limited to antigen-experienced T cells. However, we also identified populations of T cells with naïve surface phenotypes that expressed YFP; around 2% of naïve CD4^+^ cells in the spleen and colon, 16% of naïve CD8^+^ cells in the spleen and 7% of naïve CD8^+^ cells in the colon (Figs. 1C-D, Supplemental Fig. 1H). To determine whether naïve CD4^+^ T cells had previously expressed T-bet or were actively expressing T-bet, we utilised a T-bet AmCyan reporter line (57). We found that around 0.5% of naïve CD4^+^ T cells and 8% of CD4^+^ T_EM_ cells were T-bet^+^ (Figs. 1E and F). We conclude that although T-bet expression is higher in T_EM_ and TCM populations, there exist populations of CD4^+^ and CD8^+^ T cells that display evidence of T-bet expression in the context of a seemingly naïve state, which has not previously been reported.

### Naïve CD4^+^ T cells develop and exit the thymus without expressing T-bet

We sought to determine whether the expression of YFP in a proportion of naïve CD4^+^ T cells indicated prior T-bet expression in the thymus. We found that thymic double negative (DN)3, DN4, double positive (DP) and CD4 single positive (SP) cells completely lacked YFP expression (Figs. 2A and B and Supplemental Fig. 2A). Unexpectedly, the CD3^−^ CD44^+^ CD25^−^ and CD3^−^ CD44^+^ CD25^+^ populations exhibited a high proportion of YFP^+^ cells (32% and 5% respectively; Figs. 2A and B). However, further immunophenotyping demonstrated that the majority (77%) of thymic YFP^+^ cells were in fact type-1 innate lymphoid cells (ILC1s) (CD127^+^ NK1.1^+^ CD122^+^; 35% of cells; Fig. 2C, Supplemental Fig. 2B) or CD3^+^ NK cells (CD122^+^ NK1.1^+^ CD3e^+^; 42% of cells; Fig. 2C, Supplemental Fig. 2B). Other YFP-positive cells in the thymus included a small population of effector (CD62L^−^ CD44^+^) CD4^+^ T cells, of which 50% expressed YFP, and 5% of thymic T_REG_ (Figs. 2A and B). However, based on the lack of YFP expression in DN3, DN4, DP and CD4 SP cells, we conclude that all mature SP naïve CD4 cells leave the thymus YFP-negative and only express T-bet, becoming YFP-positive, in the periphery.

**Figure 2.**
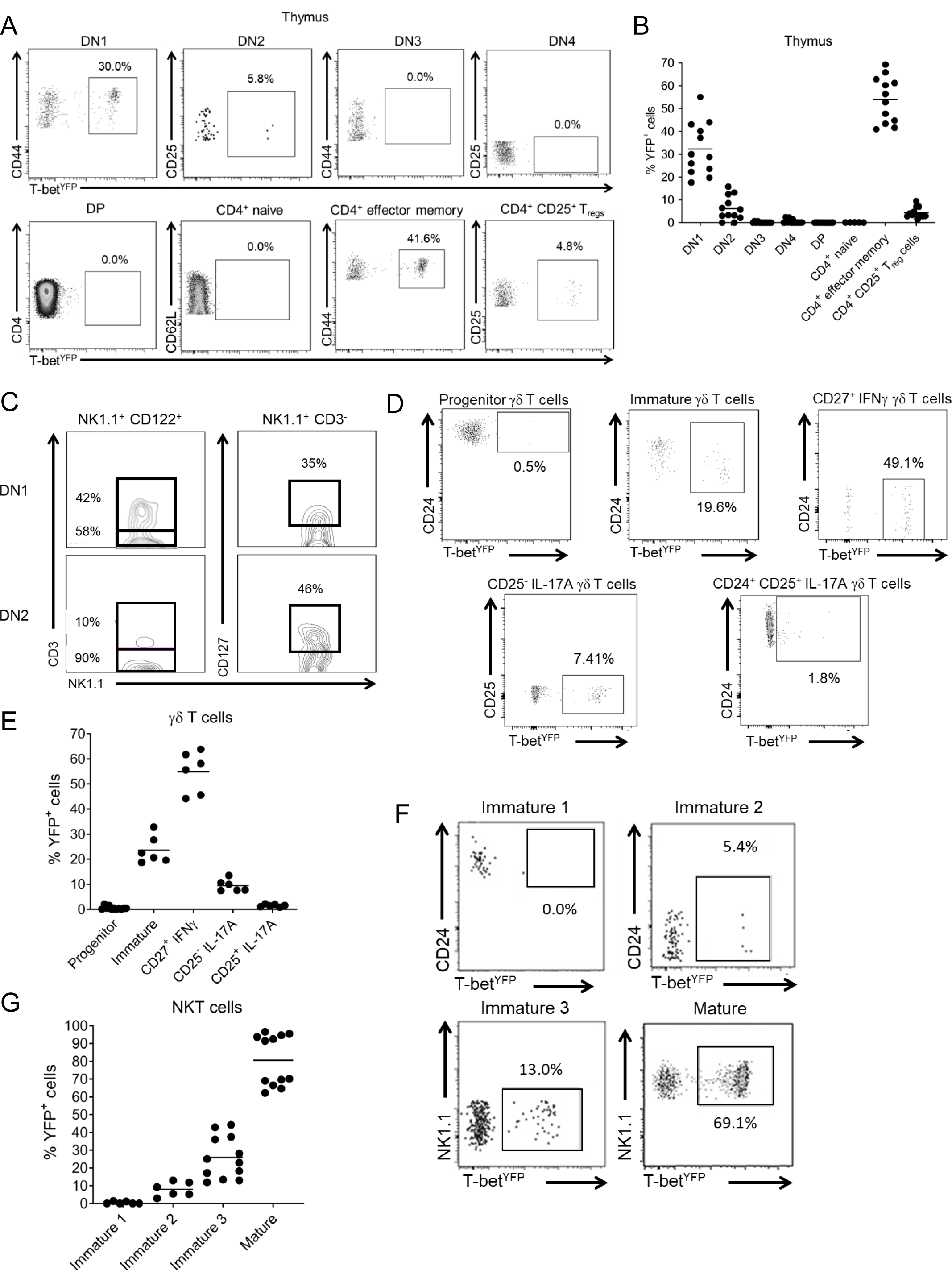
Naïve CD4^+^ T cells develop and exit the thymus without expressing T-bet. A. Representative flow plots showing YFP expression in thymic DN1 (CD3^−^ CD44^+^ CD25^−^), DN2 (CD3^−^ CD44^+^ CD25^+^), DN3 (CD3^+^ CD44^−^ CD25^+^), DN4 (CD3^+^ CD44^−^ CD25^−^), DP (CD3^+^ CD4^+^ CD25^+^), naïve CD4^+^ T cells (CD3^+^ CD4^+^ CD8^−^ CD62L^+^ CD44^−^), memory CD4^+^ T cells (CD3^+^ CD4^+^ CD8^−^ CD62L^−^ CD44^+^) and regulatory CD4^+^ T cells (CD3^+^ CD4^+^ CD8^−^ CD25^+^). B. Mean proportions of YFP^+^ cells in the different thymic cell subsets shown in A (n=12). C. Representative flow plots of NK and NKT markers on YFP^+^ cells from DN1 (live CD4^−^ CD8^−^ CD44^+^ CD25^+^ CD122^+^ NK1.1^+^) and DN2 (live CD4^−^ CD8^−^ CD44^+^ CD25^+^ CD122^+^ NK1.1^+^) cells from the thymus. D. Representative flow plots showing YFP expression in five populations of developing γδT cells in the thymus: progenitor γδT cells (CD25^+^ CD27^+^ CD24^+^), immature γδT cells (CD25^−^ CD27^+^ CD24^+^), IFNγ-producing γδT cells (CD25^−^ CD24^−^ CD27^+^) and the two IL-17A-producing γδT cells (CD25^−^ CD27^−^ CD24^−^ and CD25^+^ CD27^−^ CD24^+^). E. Mean proportions of YFP^+^ cells from the different stages of γδT cells development shown in D (n=6). F. Representative flow plots showing YFP expression in the four populations of developing NKT cells in the thymus: Immature 1 (CD44^−^ CD24^+^), immature 2 (CD44^−^ CD24^−^), immature 3 (CD44^+^ CD24^−^ NK1.1^−^) and mature NKT cells (CD44^+^ NK1.1^+^). G. Mean proportions of YFP^+^ cells from the different stages of NKT cell development shown in F (n=12).

We next examined YFP expression in γδ T cells (Supplemental Fig. 2C), which diverge from the αβ T cell development at the DN2-DN3 stages, and NKT cells (Supplemental Fig. 2D), which arise from the DP stage in the thymus. As expected, no progenitor γδ T cells expressed YFP (Figs. 2D and 2E). Surprisingly, however, 20% of CD24^+^ immature γδ T cells were YFP^+^, meaning that some of the cells have expressed T-bet despite not being fully mature (Figs. 2D and 2E). YFP expression was primarily associated with T-bet expression, with almost 50% of CD25^−^CD27^+^CD24^−^ γδ T cells, which have been previously described as IFNγ-producers, expressing YFP. Interestingly, a small population (8%) of CD25-CD27-γδ T cells, previously reported as IL-17A-producing cells, were also YFP^+^ (Figs. 2D and 2E). We conclude that as developing γδ thymocytes mature, there is an increase in the proportion that express YFP, predisposing these cells to becoming T-bet^+^ IFNγ^+^ γδ T cells.

Next, looking at NKT cell development, we found that T-bet expression increased with cell maturation; with immature stage 1 cells (CD44^−^ CD24^+^) completely lacked YFP expression and YFP positivity increased to 5-10% at immature stage 2, 15-40% at immature stage 3, to finally 60-90% YFP positivity for mature NKT cells (Figs. 2F, 2G and Supplemental Fig. 2D,).

We conclude from this analysis that conventional CD4^+^ and CD8^+^ T cells develop in the thymus without expressing YFP and therefore only express T-bet once they have left the thymus. However, NKT cells and a subpopulation of γδ T cells, which both require T-bet to develop to maturity, show an increasing proportion of YFP^+^ cells in the thymus before leaving the thymus as mature cells, again validating our T-bet^FM^ model.

### Naïve CD4^+^ T cells become YFP^+^ shortly after birth

We sought to further characterise the population of YFP^+^ naïve-like CD4^+^ cells using additional cell surface markers, expanding our analysis to the periphery and different organs within the mice after thymic development. Dividing cells into naïve (CD62L^+^ CD44^−^ CCR7^+^ CD28^+^ CD27^+^), TCM (CD62L^+^ CD44^+^ CCR7^+^ CD28^+^ CD27^−^) and T_EM_ (CD62L^−^CD44^+^ CCR7^−^ CD28^+^ CD27^−^) populations revealed that 0.5-1% of naïve CD4^+^ T cells were YFP^+^ (Fig. 3A,3B). However, 15-38% of TCM and 34-74% of T_EM_ cells expressed YFP. We conclude, therefore, that there exists a population of CD4^+^ T cells with all the hallmarks of naïve cells state that nevertheless have expressed (or possibly continue to express) T-bet.

**Figure 3.**
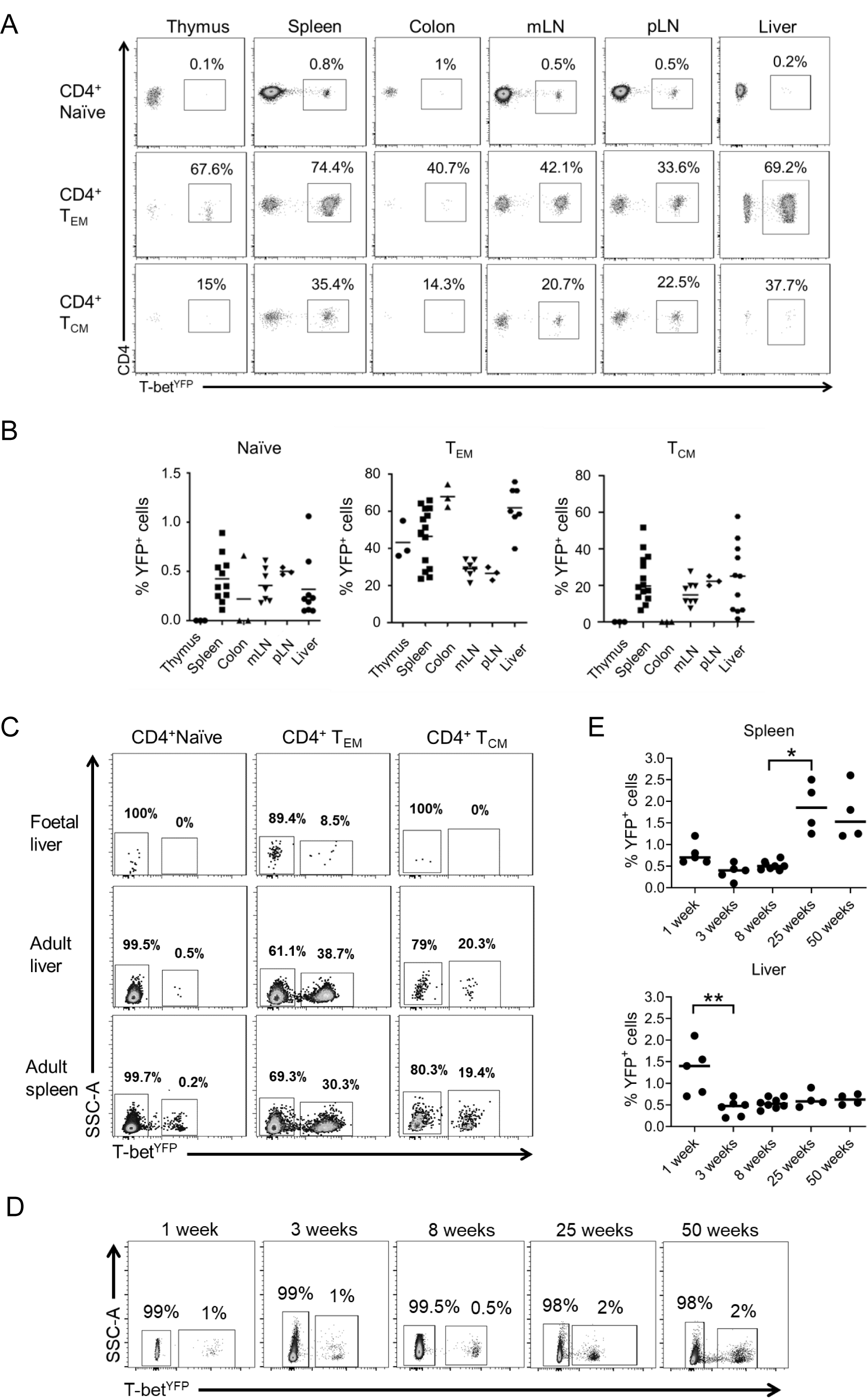
Naïve CD4^+^ T cells become YFP^+^ shortly after birth. A. Representative flow plots showing YFP expression in the naïve CD4^+^ T cells (live CD3^+^ CD4^+^ CD62L^+^ CD44^−^ CCR7^+^ CD28^+^ CD27^+^), effector CD4^+^ T cells (live CD3^+^ CD4^+^ CD62L^−^ CD44^+^ CCR7^−^ CD28^+^ CD27^−^) and central memory CD4^+^ T cells (live CD3^+^ CD4^+^ CD62L^−^ CD44^+^ CCR7^−^ CD28^−^ CD27^−^) in the thymus, spleen, colon, mesenteric lymph nodes, peripheral lymph nodes and liver. B. Mean proportions of YFP^+^ cells in naïve, effector memory and central memory CD4^+^ T cell populations shown in A (n=3 for thymus, colon, pLN, n=5 for liver, n=6 for mLN and n=8 for spleen). C. Representative flow plots showing YFP expression in naïve (CD62L^+^ CD44^−^), effector (CD62L^−^ CD44^+^) and central memory (CD62L^+^ CD44^+^) CD4^+^ T cells in the foetal liver, adult liver and adult spleen. D. Representative flow plots showing YFP expression in splenic naïve (CD62L^+^ CD44^−^) CD4^+^ T cells from 1-week (n=5), 3-week (n=6), 8-week (n=8), 25-week (n=4) and 50-week (n=4) old mice. E. Mean proportions of YFP^+^ cells in naïve (CD62L^+^ CD44^−^) CD4^+^ T cells from 1-week (n=5), 3-week (n=6), 8-week (n=8), 25-week (n=4) and 50-week (n=4) old mice. * = p<0.05, ** = p<0.01 (Kruskal-Wallis test performed with Dunn’s corrections)

We next sought to determine whether YFP^+^ naïve CD4^+^ T cells were present before birth or only arise after birth, potentially due to exposure to environmental antigens. Examination of the foetal liver at E15.5 revealed a lack of YFP^+^ naïve or TCM CD4^+^ cells, but the presence of YFP^+^ T_EM_ CD4^+^ T cells, comprising around 9% of the population (Fig. 3C). We found that by 1-week of age post-partum, 1% of naïve CD4^+^ T cells expressed YFP in the liver, and despite a significant decrease at 3 weeks of age, the proportion of YFP^+^ cells in the liver remained consistent thereon in (Fig. 3D and E). A similar pattern was observed in the spleen, with around 0.6% of cells expressing YFP in 1-week old mice, rising significantly to 1.5-2% in older animals (Fig. 3D and E). We conclude that T-bet induction in naïve-like-T cells occurs shortly after birth. This is consistent with the gain in T-bet expression being driven by the microbiota or other environmental factors.

### Naïve YFP-positive cells do not correspond to previously defined CD4^+^ T cell populations

We next sought to determine potential relationships between YFP-positive naïve CD4^+^ T cells and previously identified naïve-like memory cell populations, in particular memory T cell with naïve-like phenotype (T_MNP_ cells) (41), virtual memory (VM) T cells (37), stem cell-like memory T cells (T_SCM_) (43, 44), T_H_1-like memory phenotype (T_H_1-like MP) cells (42) and memory-phenotype (MP) cells (46).

We first measured YFP expression in T_MNP_ cells, which, like naïve cells, are CD62L^+^ CD44^-^ CD28^+^ CD27^+^, but, unlike naïve cells are also CXCR3^+^ and CD49d^+^ (41). However, we identified very few TMNP cells within the peripheral organs of T-bet^FM^ mice (Supplemental Figure 3A) and none of these cells were YFP^+^ (Fig. 4A). In contrast, 0.3% of CXCR3-CD49d-naïve CD4 T cells were YFP^+^ and 16.7% of CXCR3^+^ CD49d-cells were also YFP^+^ (Fig. 4A). This demonstrated that YFP^+^ naïve-like CD4^+^ T cells do not correspond to T_MNP_ cells. Using the same gating strategy, it was also possible to identify VM cells (CD62L^−^ CD44^+^ CD49^−^ CXCR3^+^) (37). 73% of these cells were YFP^+^, but since they were not CD62L^+^ and CD44^−^, these are distinct from the naïve YFP^+^ cell population (Fig. 4A).

**Figure 4.**
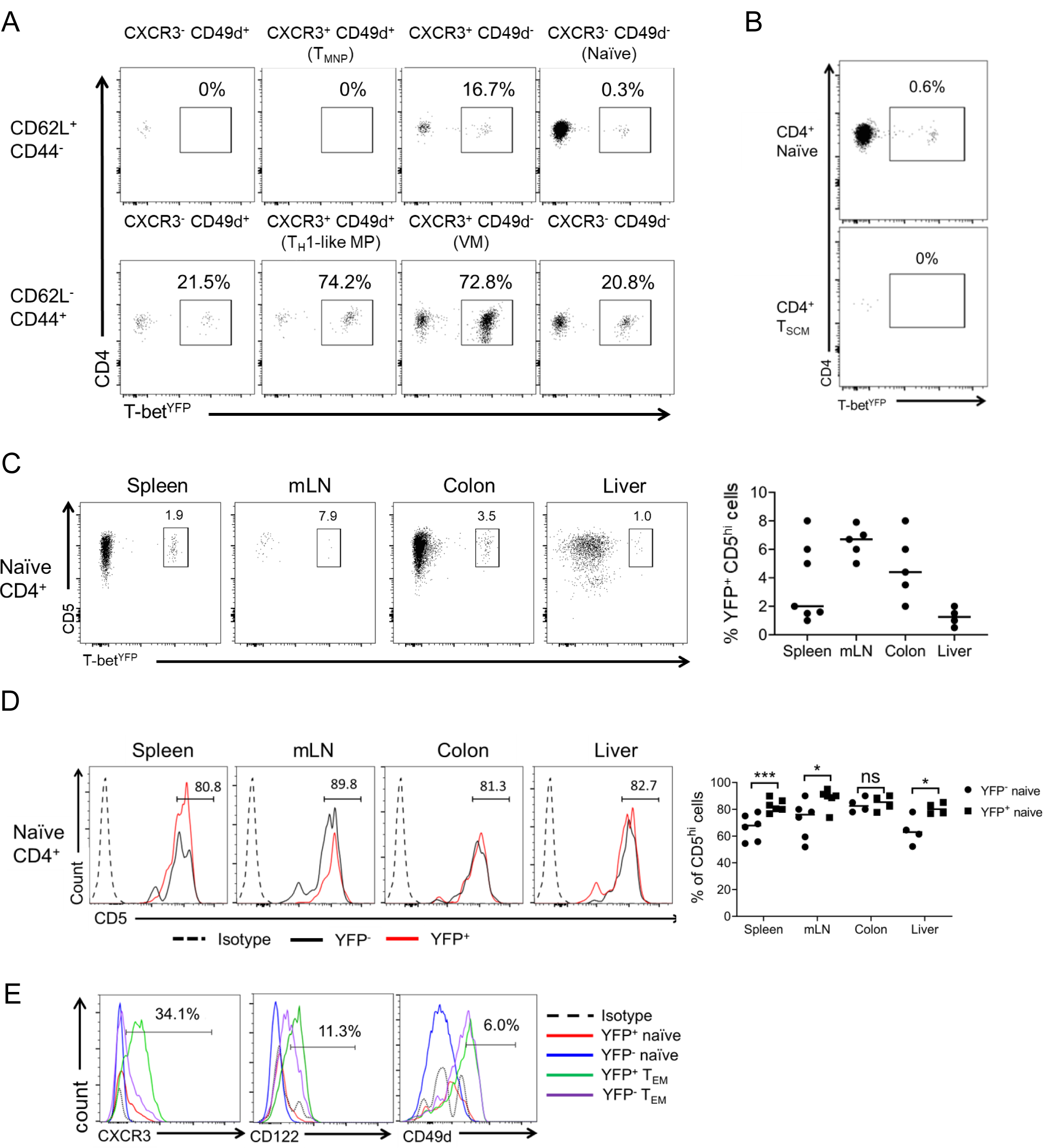
YFP^+^ naïve-like cells do not correspond to previously defined CD4 T cell populations. A. Representative flow plots showing YFP expression in splenic CD28^+^ CD27^+^ T cell populations divided by CD62L, CD44, CXCR3 and CD49d expression. Naïve (CD62L^+^ CD44^−^ CXCR3^−^ CD49d^−^), TMNP (CD62L^+^ CD44^−^ CXCR3^+^ CD49d^+^), T_H_1-like MP (CD62L^−^ CD44^+^ CD49d^+^ CXCR3^+^) and VM (CD62L^−^ CD44^+^ CD49d^−^ CXCR3^+^) T cell populations are labelled. B. Representative flow plots showing YFP expression by conventional naïve CD4^+^ T cells (CD62L^+^ CD44^−^ CD28^+^ CD27^+^ CD127^+^ CD122^−^ CD95^−^) compared with TSCM (CD62L^+^ CD44^−^ CD28^+^ CD27^+^ CD127^+^ CD122^+^ CD95^+^) from the spleen. C. Representative flow plots and mean proportions of naïve YFP^+^ CD5^hi^ CD4^+^ T cells (live CD3^+^ CD4^+^ CD62L^+^ CD44^−^) in the spleen, mLN, colon and liver (n=6 for spleen and mLN, n=5 for colon and n=4 for liver). D. Representative flow plots and mean proportions of CD5^hi^ expressing cells from naïve YFP^+^ CD4^+^ T cells (live CD3^+^ CD4^+^ CD62L^+^ CD44^−^) and naïve YFP^−^ CD4^+^ T cells in the spleen, mLN, colon and liver (n=6 for spleen and mLN, n=5 for colon and n=4 for liver). Percentages shown for YFP^+^ naïve CD4^+^ T cells. * = p<0.05, *** = p<0.005 (multiple t-test) E. Representative histograms showing surface marker expression in YFP^−^ vs YFP^+^ naïve (live CD3^+^ CD4^+^ CD62L^+^ CD44^−^) and effector memory (live CD3^+^ CD4^+^ CD62L^−^ CD44^+^) CD4^+^ T cells from the spleen. Percentages shown for YFP^+^ naïve CD4^+^ T cells.

We then measured YFP expression in T_SCM_, which like naïve cells are CD62L^+^ CD44- CD28^+^ CD27^+^ IL-7R^+^ but also express the memory cell markers CD122^+^ and CD95^+^ (43, 44). We identified very few of these cells but found that they were uniformly YFP-negative (Fig. 4B and Supplemental Fig. 3B). In contrast, 0.6% of the canonical naïve CD4^+^ population (CD62L^+^ CD44^−^ CD28^+^ CD27^+^ IL-7R^+^ CD122^−^ CD95^−^) were YFP^+^ (Fig. 4B). Thus, naïve YFP^+^ CD4^+^ T cells are distinct from T_SCM_.

Finally, we examined T_H_1-like MP cells (42) and found that 74.2% of these cells were YFP^+^ (Fig. 4A), consistent with that prior demonstration that these cell express T-bet and produce IFNγ (42). However, unlike naïve cells, T_H_1-like MP cells are CD62L^−^ CD44^+^ CXCR3^+^ CD49d^+^ (42) and thus are distinct from the naïve-like YFP^+^ cell population. Naïve YFP^+^ cells are also distinct from another described population of MP cells, which are CD62L^−^ CD44^+^ CD28^+^ CD5^hi^ (46).

CD5^hi^ naive cells have been found to generate more MP cells than their CD5^lo^ counterparts in the absence of foreign antigen, suggesting an involvement of reactivity to self-antigen in MP cell generation (46). Thus, we considered that the naïve-like YFP^+^ cell population that we observed may also be CD5^hi^. We found that only a small minority of naïve CD5^hi^ cells were YFP^+^ (Fig. 4C) but that a higher proportion of YFP^+^ cells were CD5^hi^ compared to YFP^−^ cells (Fig. 4D), consistent with exposure to self-antigen.

Lastly, we sought to investigate potential differences in the expression of the typical activated memory markers CD122, CD49d and CXCR3 between YFP^−^ and YFP^+^ CD4^+^ naïve (CD62L^+^ CD44^−^) and T_EM_ (CD62L^−^ CD44^+^) cells. We found that both YFP^−^ and YFP^+^ naïve CD4^+^ T cells exhibited lower levels of CD49d and CD122 in comparison to both YFP^+^ and YFP^−^ T_EM_ populations (Fig. 4E). This again suggests that YFP^+^ naïve cells are not a memory population. Interestingly, when comparing YFP^+^ naïve against YFP^−^ naïve T cells, the only major difference in surface marker expression was CXCR3 (a T-bet target gene), which was present on 30% of YFP^+^ naïve cells compared with only 3% of YFP^−^ naïve CD4^+^ T cells (Fig. 4E). We conclude that YFP^+^ naïve CD4^+^ T cells share naïve surface markers with YFP^−^ naïve CD4^+^ T cells except for an increased CXCR3 expression and do not represent a previously described T cell subset. Given this characterisation, we will refer to the phenotype of these cells as naïve from here on in.

### YFP^+^ naïve CD4^+^ T cells are predisposed to produce IFNγ upon activation

We hypothesised that YFP^+^ naïve cells may possess distinct properties to YFP^−^ naïve cells and that their experience of T-bet expression may allow rapid upregulation of IFNγ. To test this, we purified YFP^−^ and YFP^+^ naïve (CD62L^+^ CD44^−^) and T_EM_ (CD62L^−^ CD44^+^) CD4^+^ T cells, activated these cells *in vitro* in non-polarising conditions and measured cytokine production and T-bet expression by ELISA and flow cytometry. We found that activation of naive YFP^+^ cells caused robust induction of IFNγ that was not observed in the YFP^−^ population (Fig. 5A, Supplemental Fig. 4A). In contrast, there was no production of IL-17A, suggesting that naïve YFP^+^ cells are specifically polarised towards the T_H_1 lineage. T-bet was present in over 90% of IFNγ-producing YFP^+^ naïve cells, whereas YFP^−^ naïve cells lacked T-bet expression (Fig. 5A). We also examined the effect of stimulation on cytokine production by YFP^+^ T_EM_ cells. As we found for their naïve counterparts, YFP^+^ T_EM_ cells expressed IFNγ, but not IL-17A, whereas YFP^−^ effector cells expressed either IFNγ or IL-17A (Fig. 5B, Supplemental Fig. 4A). Further examination found that naïve YFP^+^ cells were able to rapidly upregulate IFNγ, even after 3 days of culture in vitro in non-polarising conditions, when compared to naïve YFP^−^ naïve cells (Supplemental Figs. 4B and 4C).

**Figure 5.**
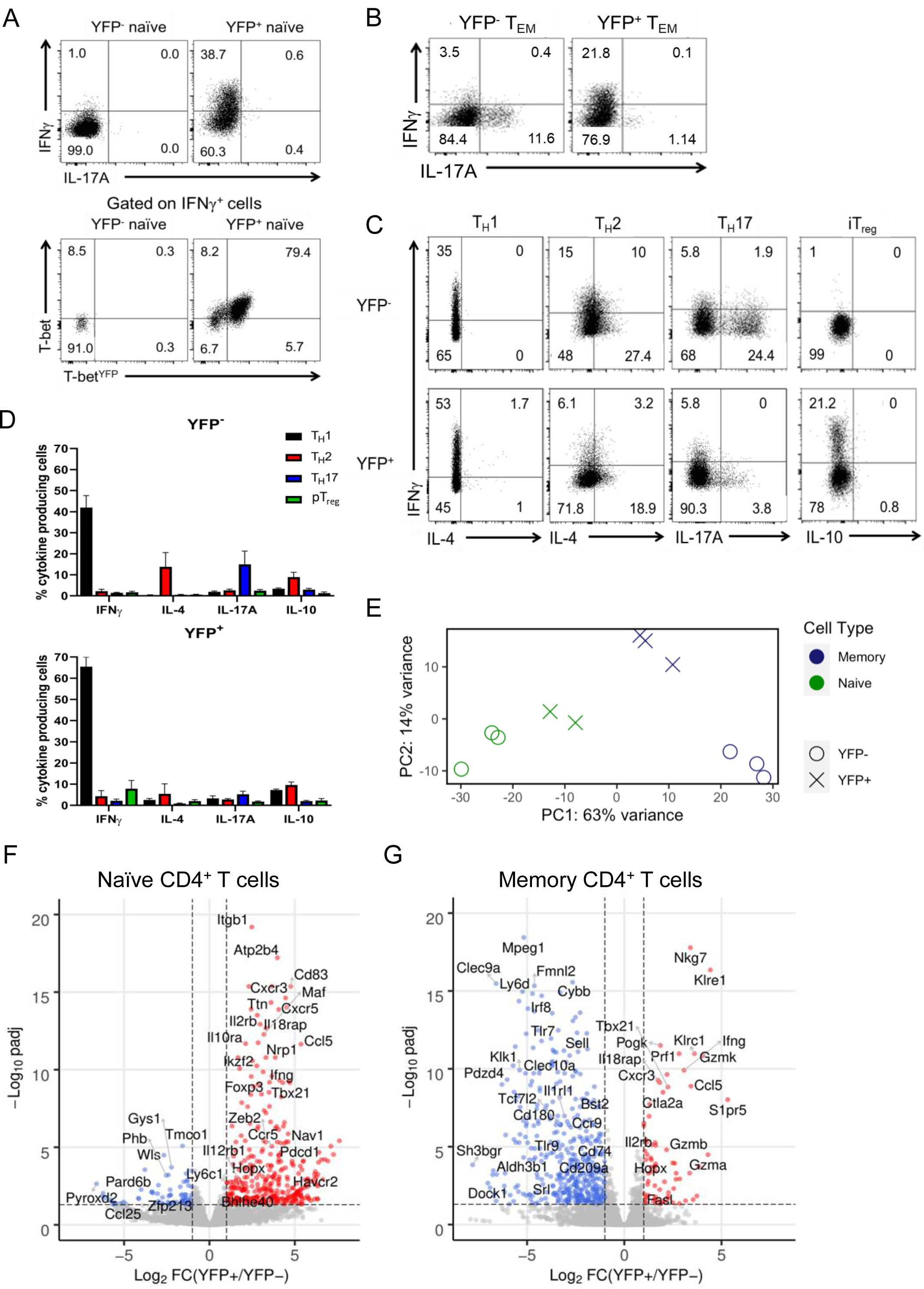
YFP^+^ naïve-like CD4^+^ T cells are polarised towards the T_H_1 lineage. A. Representative flow plots showing cytokine expression (top) or T-bet and YFP expression (bottom) in YFP^−^ and YFP^+^ naive (CD62L^+^ CD44^−^) CD4^+^ T cells after activation with anti-CD3/CD28 and culture with IL-2. B. Representative flow plots showing cytokine expression from *in vitro* cultured YFP^−^ and YFP^+^ memory (CD62L^+^ CD44^−^) CD4^+^ T cells after activation with anti-CD3/CD28 and culture with IL-2. C. Representative flow plots showing cytokine expression in naïve (CD62L^+^ CD44^−^) CD4^+^ T cells after activation with anti-CD3/CD28 activation and polarisation in T_H_1, T_H_2 T_H_17 and iTREG conditions *in vitro* (n=6). D. Cytokine expression (mean and SEM (n=6)) in naïve (CD62L^+^ CD44^−^) CD4^+^ T cells after activation with anti-CD3/CD28 and polarisation in T_H_1, T_H_2 T_H_17 and iTREG conditions *in vitro*. E. PCA plot showing the relationship between the gene expression profiles of YFP^+^ and YFP^−^ naïve (live CD3^+^ CD4^+^ CD62L^+^ CD44^−^) and T_EM_ (live CD3^+^ CD4^+^ CD62L^−^ CD44^+^) CD4^+^ T cell populations. F. Volcano plot showing differential gene expression between YFP^+^ and YFP^−^ naïve CD4^+^ T cells (log2 fold change vs adjusted p-value). Genes more highly expressed in YFP^+^ cells are in red and those more highly expressed in YFP^−^ cells are in blue. Selected differentially expressed genes are labelled. G. Volcano plot showing differential gene expression between YFP^+^ and YFP^−^ T_EM_ CD4^+^ T cells (log2 fold change vs adjusted p-value). Genes more highly expressed in YFP^+^ cells are in red and those more highly expressed in YFP^−^ cells are in blue (p_adj_ <0.05, |log2FC|>1). Selected differentially expressed genes are labelled.

We next sought to determine whether the predisposition of YFP^+^ naïve cells to produce IFNγ was stable under T_H_2, T_H_17 and iT_reg_ polarising conditions. The naïve YFP^+^ population resisted induction of IL-4 or IL-17A after culture in T_H_2 or T_H_17-polarising conditions, respectively (Figs. 5C and 5D). Similarly, compared to YFP^−^ naïve cells, a greater proportion of YFP^+^ naïve cells maintained IFNγ production under T_H_2 and iTREG-polarising conditions, although induction was weaker compare to T_H_1 conditions (Fig. 5C and 5D). Staining for lineage-specific transcription factors confirmed that YFP^+^ cells resisted re-polarisation to other lineages (Supplemental Fig. 4D). We conclude that naïve YFP^+^ cells are predisposed to produce IFNγ and express T-bet over cytokines and transcription factors of other T cell lineages when polarised *in vitro*.

### Naïve and T_EM_ YFP^+^ CD4^+^ T cells are polarised towards a T_H_1 phenotype

We wished to determine whether the predisposition of YFP^+^ naïve cells to produce IFNγ reflected polarisation towards the T_H_1 lineage. To test this, we compared the gene expression profiles of YFP^+^ versus YFP^−^ naïve and CD62L^−^ CD44^+^ T_EM_ CD4^+^ T cells using RNA-seq. Visualising the relationship between the expression profiles of the four cell populations, we found that YFP^+^ naïve and YFP^+^ T_EM_ cells were distinct from their YFP^−^ counterparts, with YFP^+^ naïve cells being more similar to YFP^−^ naïve cells than to either of the T_EM_ cell populations (Fig. 5E and Supplemental Fig. 4E). This is consistent with our flow cytometric analyses showing that YFP^+^ naïve cells display the hallmarks of a naïve cell population.

We next sought to identify the genes that distinguished YFP^+^ naïve cells from YFP^−^ naïve cells. We found that YFP^+^ cells exhibited high levels of expression of a number of T-bet target genes that mark T_H_1 cells, including *Tbx21, Ifng, Cxcr3, Ccl4, Ccl5, Itgb1, Havcr2, Nkg7, Il12rb1* and *Il18rap*, and genes encoding transcription factors that cooperate with T-bet, including *Zeb2, Hopx* and *Bhlhe40* (Fig. 5F). Consistent with this, gene set enrichment analysis (GSEA) confirmed enrichment of T_H_1 genes within the set of genes highly expressed in YFP^+^ versus YFP^−^ naïve cells (Supplemental. Fig. 4F). Thus, these data demonstrate that YFP^+^ naïve cells are polarised towards a T_H_1 lineage phenotype. We also noted higher expression of *Foxp3*, *Ikzf2* (Helios) and *Nrp1*, required for T_REG_ function, suggesting there might also be heterogeneity within the YFP^+^ naïve cell population.

We next turned our attention to the expression profile of YFP^+^ T_EM_ cells compared to their YFP^−^ counterparts. As for the naïve cells, we identified higher expression of several T_H_1 genes, including *Ifng, Tbx21, Cxcr3, Ccl5, Il12rb2, Il18rap, Nkg7, Prf1 and Fasl* (Fig. 5G). YFP^+^ T_EM_ cells also exhibited reduced expression of a broad set of genes suggesting that in this differentiated population, the repression of alternative differentiation pathways is a key function of T-bet expression. We conclude that YFP^+^ naïve and memory subsets have distinct gene expression profiles from their YFP^−^ counterparts, which is consistent with T-bet function in these cells.

### Naïve YFP^+^ CD4^+^ T cells remain T_H_1 polarised in a murine T_H_1/T_H_17 model of colitis

The role of T_H_1 and T_H_17 cells in inflammatory disease is reflected by the induction of colitis upon adoptive transfer of naïve CD4^+^ T cells into *Rag2*^*−/−*^ mice, which is marked by a wasting phenotype, inflamed colons, splenomegaly and a T_H_1, T_H_17 and T_H_1/T_H_17 dual/hybrid-driven inflammatory response (27). To test whether the T_H_1 polarised phenotype of YFP^+^ naïve cells altered their ability to induce inflammatory disease, we transferred either 25,000 YFP^−^ or YFP^+^ naïve CD4^+^ T cells into separate *Rag2*^*−/−*^ mice and observed disease symptoms and cytokine production in colon organ cultures and in unfractionated cells from spleen, mLN and colon by ELISA. Neither group of mice developed wasting disease, most likely due to the small number of naïve T cells available for transfer (the typical model involves transfer of 500,000 naïve CD4^+^ T cells). However, both sets of recipient mice gained significantly less weight compared to control mice (Fig. 6A). Next, examining colon and spleen weights, we found that mice receiving YFP^+^ naïve T cells had smaller colons (p<0.05) and spleens in comparison with YFP^−^ naïve T cell recipient mice (Fig. 6B), suggesting that YFP^+^ CD4^+^ T cells induced less inflammation than their YFP^−^ counterparts. Donor CD4^+^ T cells could be detected in the colon, mLN and spleen after 9 weeks of transfer. Measuring cytokine production by donor cells in these organs, we found no difference in the proportion of IFNγ-producing CD4^+^ T cells in YFP^+^ versus YFP^−^ treated mice (Figs. 6C and D). There was a decrease in IL-17A producers in the colon from YFP^+^ transferred mice, although this was not significant. However, the level of IFNγ production measured by ELISA in the supernatant from colonic biopsy organ culture (Fig. 6E) and from bulk cultured cells (Fig. 6F) from the colon from YFP^+^ naïve T cell recipient mice was significantly greater than for YFP^−^ naïve T cell recipients in comparison to the mice that received no transfer at all. This demonstrates that either the predisposition of YFP^+^ CD4^+^ T cells to produce IFNγ is maintained *in vivo* or that receipt of YFP^+^ CD4^+^ T cells increases IFNγ production by other cells in the mice.

**Figure 6.**
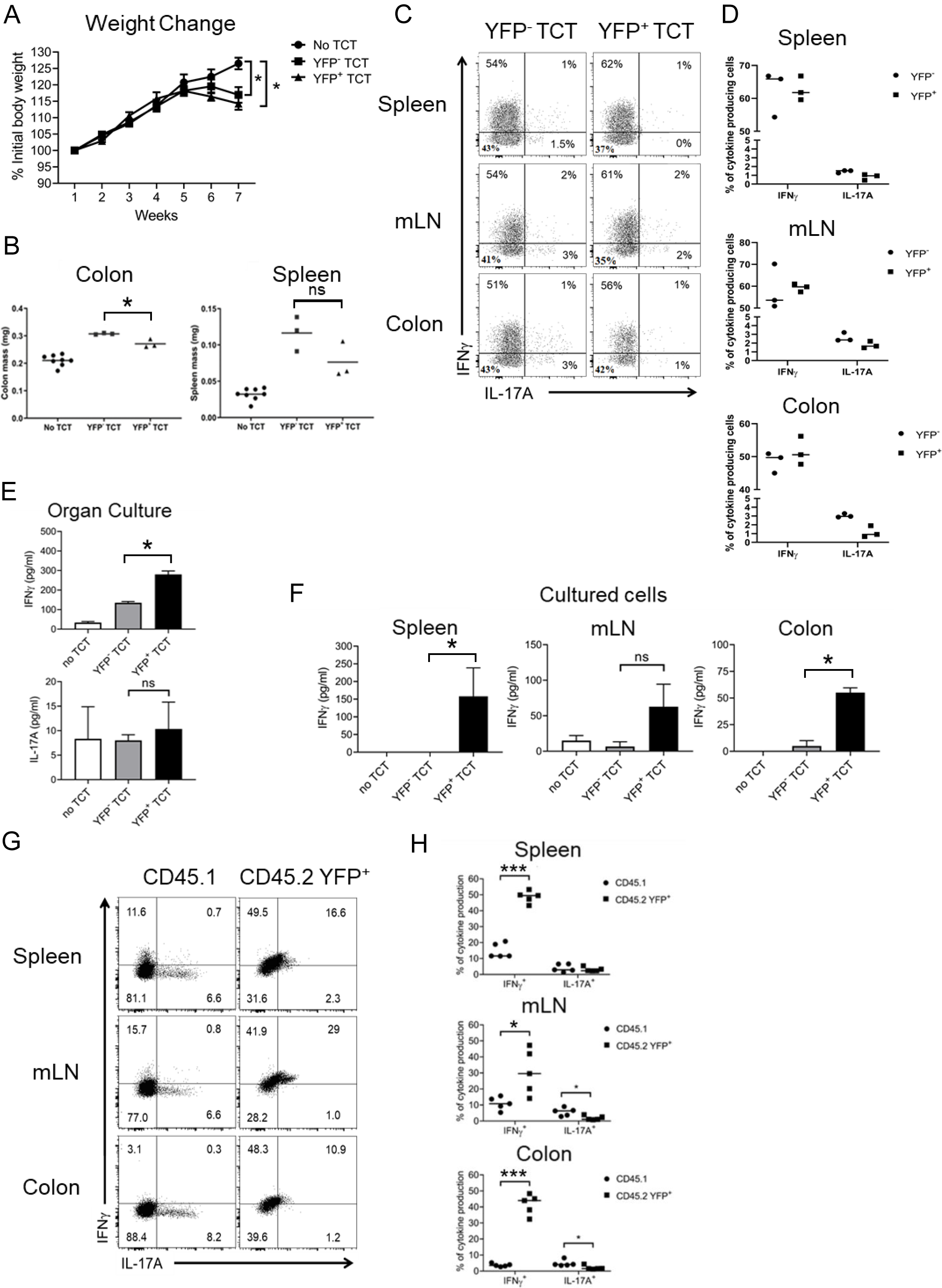
YFP^+^ cells induce less inflammation but more IFNγ compared to conventional naïve CD4^+^ T cells after transfer to *Rag2*^*−/−*^ mice. A. Weight change in *Rag2*^*−/−*^ mice after receipt of no T cells or 25,000 purified naïve YFP^+^ or naïve YFP^−^ CD4^+^ T cells (n=3 for each transfer and n=7 for control) * p<0.05 (multiple t-test). B. Spleen and colon mass in *Rag2*^*−/−*^ mice after receipt of no T cells or 25,000 purified naïve YFP^+^ or naïve YFP^−^ CD4^+^ T cells (n=3 for each transfer and n=7 for control) * = p<0.05. (Kruskal-Wallis test performed with Dunn’s corrections) C. Representative flow plots showing the proportion of YFP^−^ and YFP^+^ naïve CD4^+^ T cells that produce IFNγ and IL-17A after transfer into *Rag2*^*−/−*^ mice, separated by organ. D. Mean proportion of YFP^−^ and YFP^+^ naïve CD4^+^ T cells shown in C that produce IFNγ and IL-17A after transfer into *Rag2*^*−/−*^ mice, separated by organ (n=3 for each transfer). E. Quantification of IFNγ and IL-17A in the supernatant by ELISA after 48 hours of colon organ culture (n= 3, except n=6 for the no transfer control). * = p<0.05 (Kruskal-Wallis test performed with Dunn’s corrections) F. Quantification of IFNγ and IL-17A in the supernatant of unfractionated cell cultures from colon, spleen and mLN (n= 3, except n=6 for the no transfer control). * = p<0.05 (Kruskal-Wallis test performed with Dunn’s corrections) G. Mouse weights after co-transfer of 50,000 naïve (CD62L^+^ CD44^−^) CD45.1 and CD45.2 YFP^+^ CD4^+^ T cells at a ratio of 90:10 in *Rag2*^*−/−*^ mice (n=5). H. Representative flow plots showing IFNγ and IL-17A production by naïve CD45.1^+^ and CD45.2 YFP^+^ CD4^+^ T cells after co-transfer 90:10. I. Overall statistical dot plot showing IFNγ and IL-17A production by naïve CD45.1^+^ and CD45.2 YFP^+^ CD4^+^ T cells after co-transfer 90:10. * = p<0.05, *** = p<0.001 (multiple t-test).

In order to test whether YFP^+^ naïve CD4^+^ T cells were remained committed to producing IFNγ even in the T_H_17-polarising environment of the T cell transfer colitis model, we co-transferred naïve CD45.2 YFP^+^ CD4^+^ T cells and naïve CD45.1 CD4^+^ T cells (total of 50,000 cells at a ratio of 1:9) into *Rag2*^*−/−*^ mice. The transferred YFP^+^ CD45.2 population contained significantly more IFNγ-positive cells and lacked IL-17A-positive cells, as expected, compared to CD45.1 cells, which were able to produce IL-17A (Figs. 6G and 6H). We conclude that YFP^+^ naïve CD4^+^ T cells remain polarised to the T_H_1 lineage, even in a T_H_1/T_H_17-driven colitis model *in vivo*.

## Discussion

Here, we report the use of a novel T-bet^FM^ mouse line to identify cells that have expressed the T_H_1 lineage-specifying transcription factor T-bet. The T-bet^FM^ mouse line has allowed confirmation of known T-bet expressing lineages and revealed previously uncharacterised T-bet expressing cell populations. We identify a distinct sub-population of CD4^+^ T cells that have a naïve surface phenotype but that have experienced T-bet expression. The cells arise after birth, exhibit a T_H_1 expression profile, and rapidly upregulate IFNγ upon activation. The T_H_1 phenotype is stable both in repolarising conditions in vitro and in a T_H_1/T_H_17-driven inflammatory disease model in vivo. We conclude that T-bet expression defines a population of T_H_1-polarised naïve-like cells that function to shape subsequent T helper cell responses.

The pattern of YFP positivity, marking cells that have experienced T-bet expression, recapitulates previous knowledge of T-bet function. For example, the expression of T-bet in developing NKT cells and γδ T cells in the thymus (58–61). It has previously been reported that IFNγ-producing γδ T cells express T-bet and IL-17A producing γδ T cells express RORγt (60). However, our data suggests that within the thymus the subset of developing γδ T cells, which will develop to become IL-17A producing γδ T cells, have also expressed T-bet. Further phenotyping of these γδ T cells would be required to identify the different Vγ variable regions within these subsets (62). As expected, developing thymic SP CD4^+^ and CD8^+^ T cells and DP cells were all YFP negative. All cell types in the periphery that were expected to express T-bet-NK cells, CD4^+^ T_EM_ and T_CM_ and CD8^+^ T_EM_ and T_CM_-were positive for YFP (2, 3, 63).

Two T-bet fate mapping lines have previously been described; a T-bet-ZsGreen-T2A-CreERT2 × Rosa26-loxP-STOP-loxP-tdTomato line (57, 64) and a Tbx21tdTomato-T2A-CreERT2 × Rosa26-loxP-STOP-loxP-YFP line (65). Fang and colleagues (64) used the Tbet-ZsGreen-T2A-CreERT2 × Rosa26-loxP-STOP-loxP-tdTomato line to examine evidence for past or present T-bet expression in naïve CD4^+^ T cells but found the cells to be homogenously negative. Thus, our detection of a YFP^+^ population of naïve-like CD4^+^ T cells is in contrast with the results of this study. The lack of evidence for T-bet expression in this previous study could be due to the use of a Tmx-inducible Cre construct that only measures T-bet gene activity in the window in which Tmx is administered, as opposed to the Cre construct we use here that only depends on T-bet expression. Importantly, we could also detect T-bet expression in a subset of naïve CD4^+^ T cells using T-bet^Amcyan^ reporter mice. Furthermore, RNA-seq also demonstrated high expression of *Tbx21* in YFP^+^ versus YFP^−^ naïve T cells. Thus, these independent measurements provide additional evidence that T-bet is expressed by a population of naïve-like CD4^+^ T cells. T-bet-expressing naïve-like CD4^+^ T cells were not detected in a previous study using T-bet-ZsGreen reporter mice (64); this difference in results compared to the T-bet reporter data shown here could be due to differences in BAC integration or the T-bet construct used (T-bet-AmCyan vs T-bet-ZsGreen-T2A-CreERT2).

Characterisation of YFP^+^ naïve-like CD4^+^ T cells revealed them to lack the markers of T_SCM_, T_MNP_, T_H_1-like MP or VM cells (37, 41–43, 66, 67) and thus, we suggest that these cells represent a previously unappreciated distinct population. YFP^+^ naïve CD4 cells were CD5^hi^ and, since CD5^hi^ naïve cells have previously been shown to generate more MP cells than their CD5^lo^ counterparts (46), we suggest that YFP^+^ naïve cells may be precursors to T-bet^hi^ MP cells. However, experiments tracing the ontogeny of MP cells will be required to confirm this. Interestingly, 30% of the YFP^+^ naïve CD4 T cells were positive for CXCR3. CXCR3, a known T-bet target gene, is required for the migration of both CD4^+^ and CD8^+^ T cells to sites of inflammation (5, 56, 68, 69). Thus, YFP^+^ CXCR3^+^ CD4^+^ T cells could be predetermined early immune responders that are able to migrate to areas of infection where they can rapidly induce a T_H_1 response.

We found that YFP^+^ cells are absent from foetuses and only develop after birth, when they become visible from as early as 1 week of age and remain throughout the lifespan of the mice. This suggests a requirement for an environmental antigen in the induction of T-bet expression in these cells. This further supports the idea that T-bet^+^ naïve CD4^+^ T cells are early T_H_1-polarised inflammatory responders that react to environmental antigens.

*In vitro* and *in vivo* analyses of YFP^+^ naïve T cells show that they have a different phenotype to their YFP^−^ counterparts. The cells exhibit a T_H_1 expression profile and produce high amounts of IFNγ upon activation. YFP^+^ naïve T cells also resist polarisation to different lineages both *in vitro* and in a T_H_1/T_H_17-driven colitis model *in vivo*. These data are again consistent with a role for these cells as early T_H_1-polarised responders whose role is to produce IFNγ within inflammatory environments. These data also suggest that the YFP^+^ naïve T cells produce a more limited colitogenic phenotype due to their lack of IL-17A production, which is required for induction of colitis in the naïve T cell transfer model (15, 24, 26–28). These data further emphasised that YFP^+^ naïve CD4 T cells are committed to produce IFNγ, even in a T_H_17-inducing inflammatory environment.

The expression of T-bet in cells that otherwise appear to be naïve suggests that lineage specification can proceed in the absence of overt T cell activation. Identification of signals responsible for T-bet upregulation, and why these signals do not result in the expression of canonical markers of T cell activation, will require further investigation. The use of fate mapper mouse models to study the expression or activation of other T cell lineage-specifying factors will also reveal whether this is a more general phenomenon.

In conclusion, we have identified a CD4^+^ T cell subset with a naïve surface phenotype that has experienced T-bet expression, is pre-polarised to the T_H_1 lineage, and which provides a rapid and stable source of IFNγ upon activation. This has implications for our understanding of the mechanisms that shape T helper responses in infection and inflammatory disease.

## Acknowledgements

Research was also supported by the BRC Flow Core facility at Guy’s and St Thomas’ NHS Foundation Trust and UCL. RNA-seq was performed by Paola Niola at UCL Genomics.

## Author Contributions

Study concept and design (JWL, GML, RGJ) data acquisition (JWL, MVdM, LBR), data analysis and interpretation (JWL, MVdM, SH, LBR, JFN, NGM, ES), technical support (LBR, NGM, AH, JFN, ES, IJ), obtaining funding (RGJ, GML, JKH, JFN), writing the manuscript (JWL, MVdM, RGJ), study supervision (RGJ, GML).

## Data Availability

RNA-seq data have been deposited in the Gene Expression Omnibus (GEO) with accession code GSE153805. This can be accessed with the secure token enufyogsxrutzut.

