## Supplemental Figures for "A population of CD4^+^ T cells with a naïve phenotype stably polarized to the T_H_1 lineage"

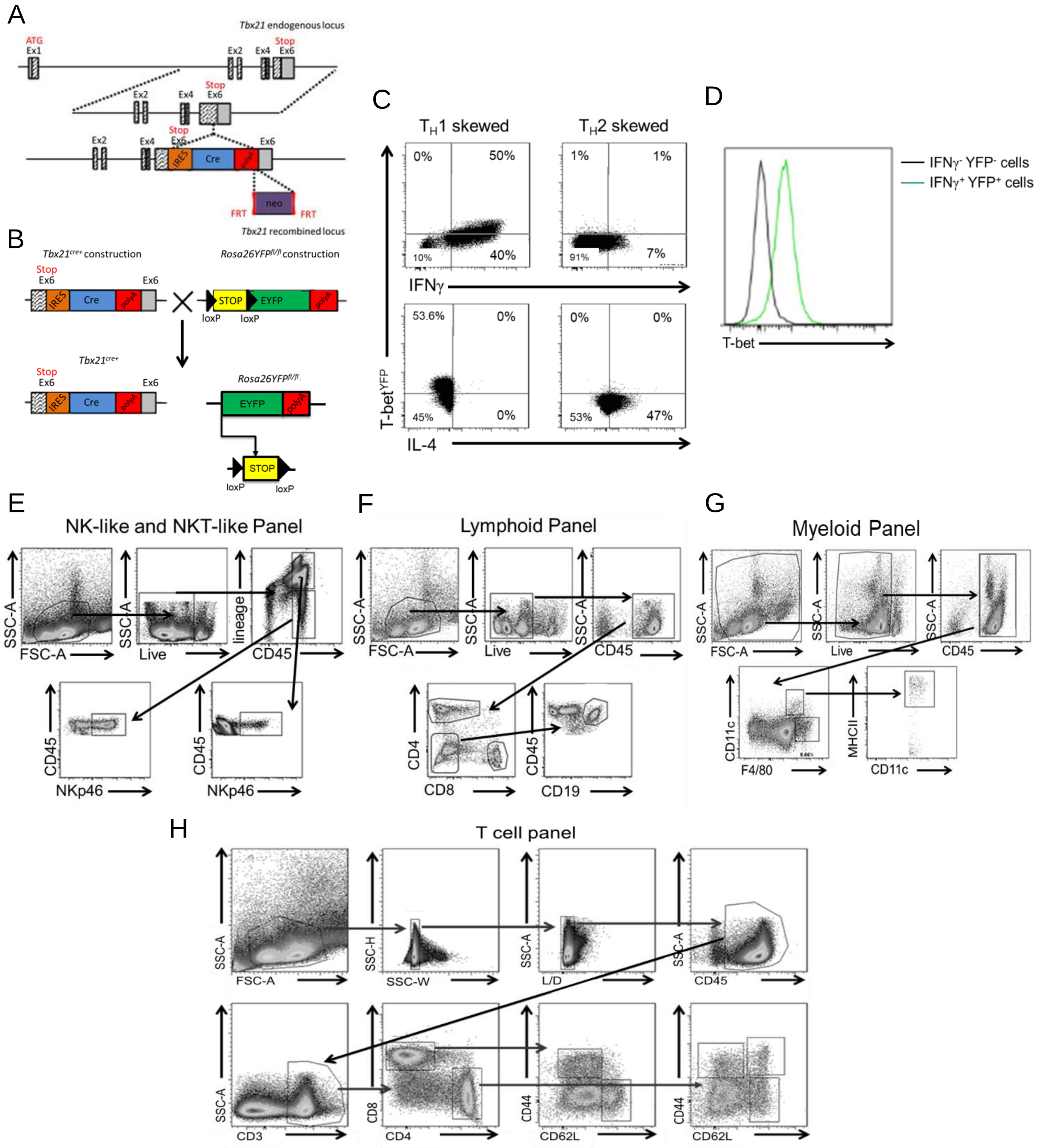


**Supplemental Figure 1. Characterising YFP expression in the T-bet$^{FM}$ mouse.**

A. Schematic showing the insertion of *Cre* into exon 6 of *Tbx21*.

B. Schematic showing the generation of the T-bet$^{cre}$ x ROSA26YFP$^{fl/fl}$ mouse line.

C. Representative flow plots showing cytokine and YFP expression in naïve (CD62L$^+$ CD44$^-$) CD4$^+$ T cells after activation with anti-CD3/CD28 activation and polarisation in T$_H$1 and T$_H$2 conditions.

D. Representative histogram showing T-bet expression of IFNγ$^+$ YFP$^+$ and IFNγ$^-$ YFP$^-$ cells.

E. Representative flow plots showing the gating strategy for NK-like and NKT-like cells.

F. Representative flow plots showing the gating strategy for lymphoid cells.

G. Representative flow plots showing the gating strategy for myeloid cells.

H. Representative flow plots showing the gating for CD4$^+$ and CD8$^+$ T cell subsets in the spleen and colon.

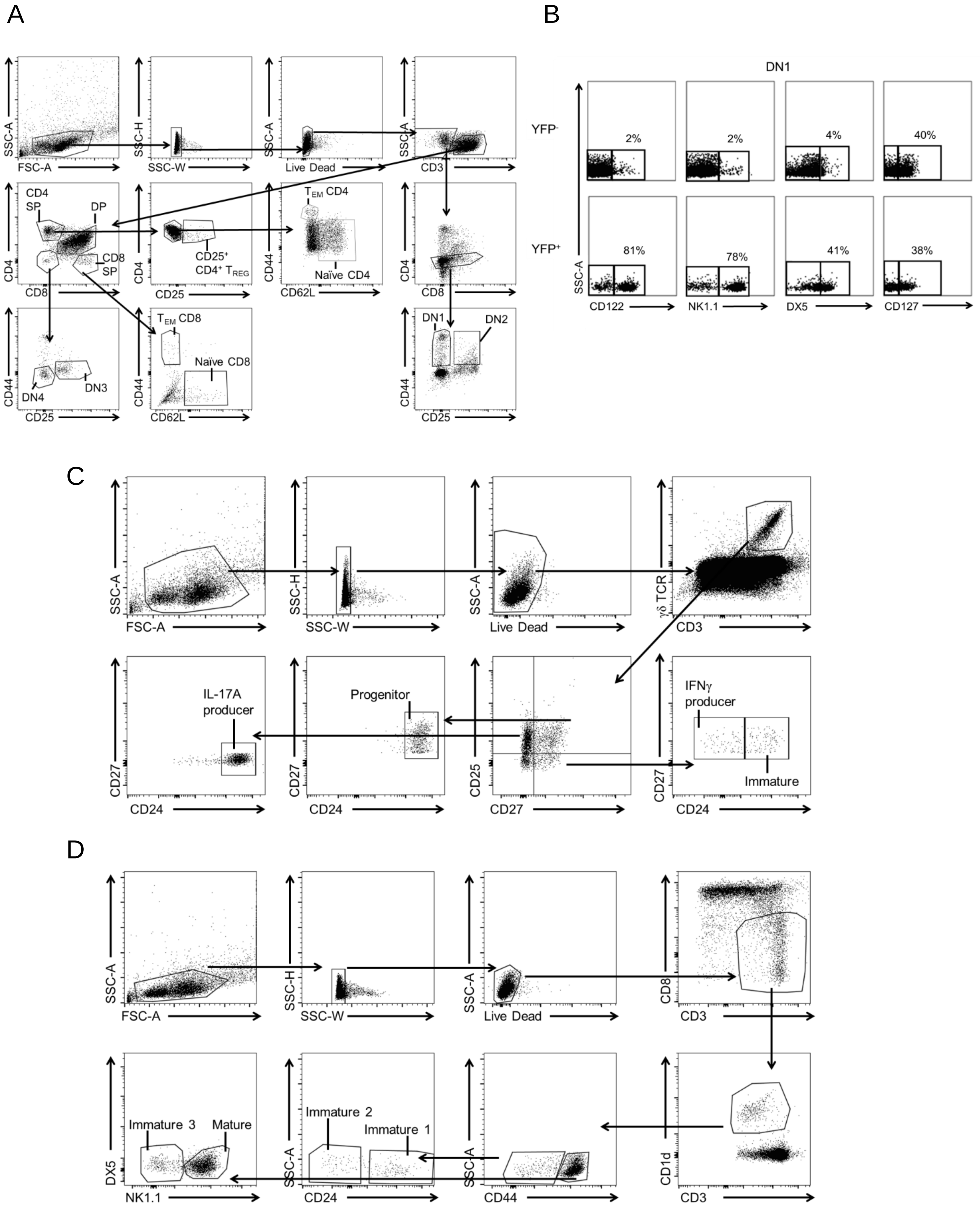


**Supplemental Figure 2. YFP expression in cells of the thymus.**

A. Representative flow plots showing the gating strategy used for cells in the thymus.

B. Representative flow plots showing the proportion of YFP⁻ and YFP⁺ DN1 cells expressing NK and NKT markers.

C. Representative flow plots showing the gating strategy used to identify γδT cells in the thymus.

D. Representative flow plots showing the gating strategy used to identify NKT cell in the thymus.

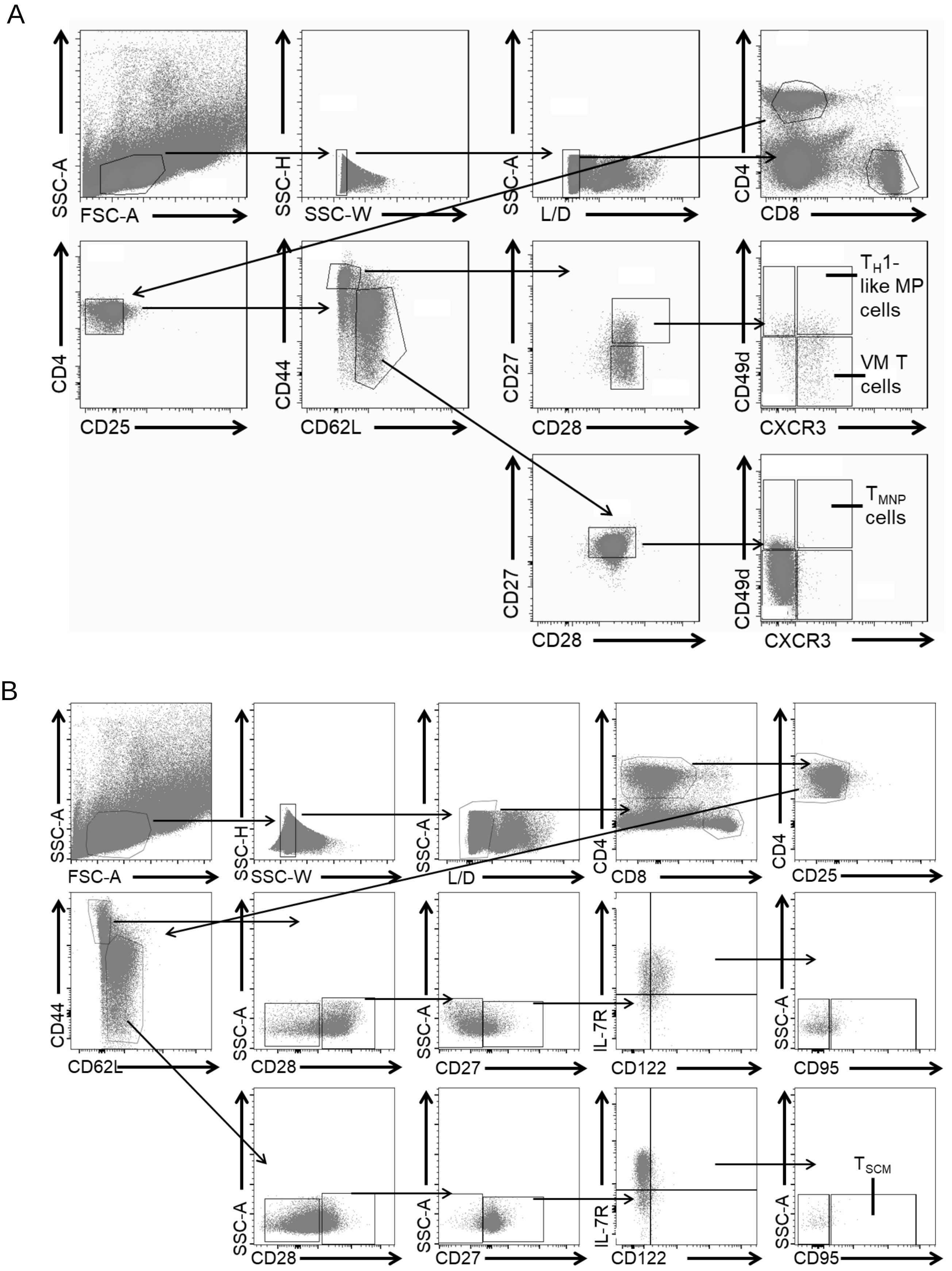


**Supplemental Figure 3. Gating strategies for $T_{SCM}$, VM, $T_H$1-like MP and $T_{MNP}$ $CD4^+$ T cells.**
A. Gating strategy shown for identifying $T_{MNP}$, $T_H$1-like MP and VM T cells in the naïve ($CD62L^+$ $CD44^-$) or memory ($CD44^+$ $CD62L^-$) compartment in the spleen.
B. Gating strategy shown for identifying $T_{SCM}$ in the naïve ($CD62L^+$ $CD44^-$) compartment in the spleen.

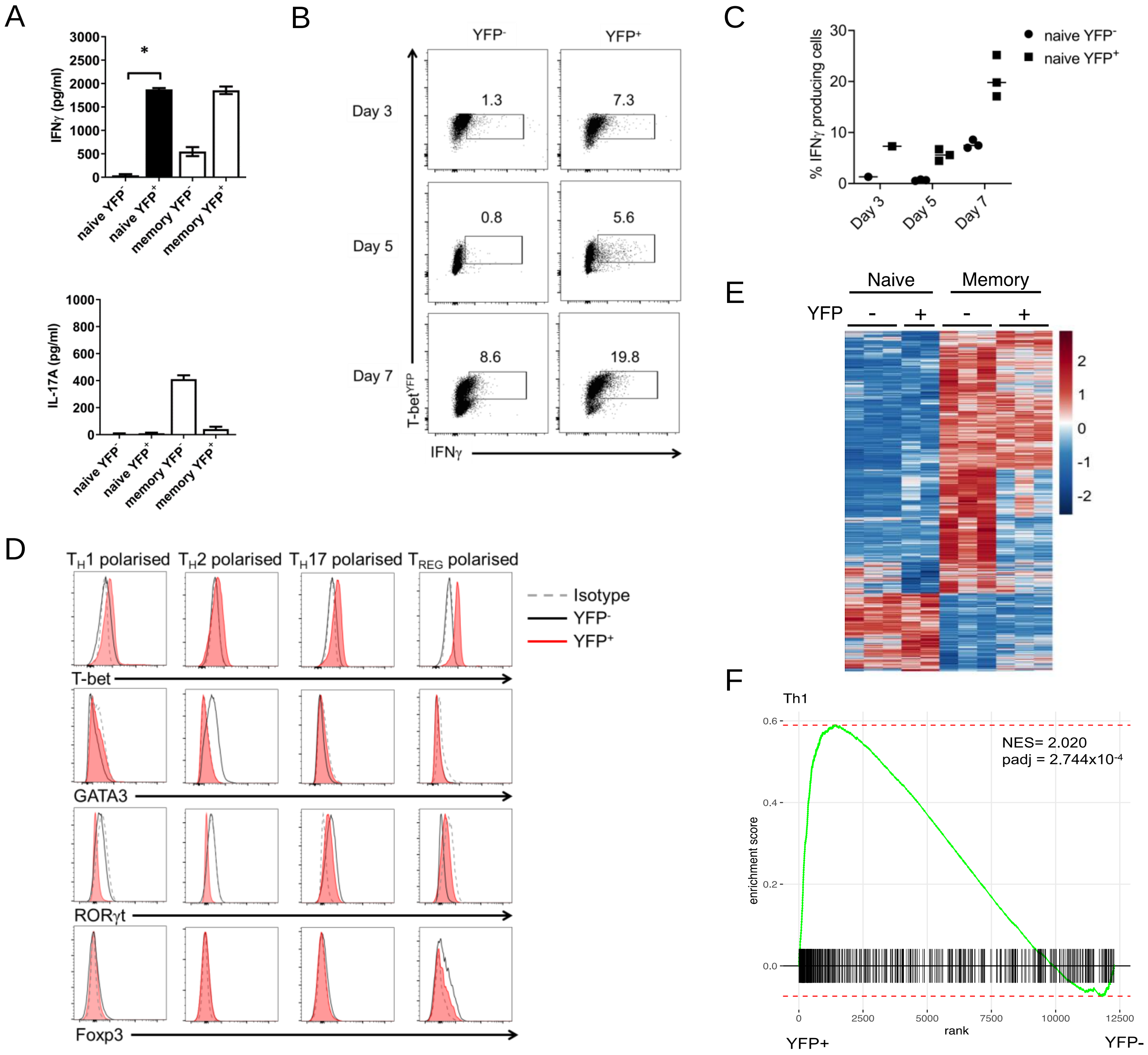


**Supplemental Figure 4. YFP$^{+}$ naïve-like CD4$^{+}$ T cells are predisposed to produce IFNγ upon activation.**

A. IFNγ and IL-17A in the supernatant of the cultured cells measured by ELISA (n=6, except n=2 for YFP$^{+}$ naïve CD4 T cells). * = P<0.05 (Kruskal-Wallis test performed with Dunn's corrections).

B. Representative flow plots showing YFP and IFNγ expression by naïve (CD62L$^{+}$ CD44$^{-}$) YFP$^{-}$ and YFP$^{+}$ CD4$^{+}$ T cells after activation with anti-CD3/CD28 and culture with IL-2 for 3, 5 and 7 days.

C. Mean proportions of naïve (CD62L$^{+}$ CD44$^{-}$) CD4$^{+}$ T cells shown in B that produce IFNγ after 3, 5 and 7 days of culture (n=3 for day 5 and 7 and n=1 for day 3).

D. Representative histograms showing transcription factor expression in naïve CD4$^{+}$ T cells after polarisation into different lineages (polarisation experiments performed three times with cells plated in triplicate for each culture).

E. Heatmap of relative gene expression between sorted naïve (CD62L$^{+}$ CD44$^{-}$) CD4$^{+}$ YFP$^{-}$, naïve CD4$^{+}$ YFP$^{+}$, effector memory (CD62L$^{-}$ CD44$^{+}$) CD4$^{+}$ YFP$^{-}$ and effector memory CD4$^{+}$ YFP$^{+}$ (n=3). Each row represents a gene and each column a cell sample. Gene expression is shown as z-score and coloured according to the scale on the right.

F. Gene set enrichment analysis (GSEA) demonstrating enrichment of $T_H1$ genes within the naïve YFP$^{+}$ CD4$^{+}$ cells versus naïve YFP$^{-}$ CD4$^{+}$ cells.
